# ARBRE: Computational resource to predict pathways towards industrially important aromatic compounds

**DOI:** 10.1101/2021.12.06.471405

**Authors:** Anastasia Sveshnikova, Homa MohammadiPeyhani, Vassily Hatzimanikatis

## Abstract

Synthetic biology and metabolic engineering rely on computational search tools for predictions of novel biosynthetic pathways to industrially important compounds, many of which are derived from aromatic amino acids. Pathway search tools vary in their scope of covered reactions and compounds, as well as in metrics for ranking and evaluation. In this work, we present a new computational resource called ARBRE: Aromatic compounds RetroBiosynthesis Repository and Explorer. It consists of a comprehensive biochemical reaction network centered around aromatic amino acid biosynthesis and a computational toolbox for navigating this network. ARBRE encompasses over 28’000 known and 100’000 novel reactions predicted with generalized enzymatic reactions rules and over 70’000 compounds, of which 22’000 are known to biochemical databases and 48’000 only to PubChem. Over 1,000 molecules that were solely part of the PubChem database before and were previously impossible to integrate into a biochemical network are included into the ARBRE reaction network by assigning enzymatic reactions. ARBRE can be applied for pathway search, enzyme annotation, pathway ranking, visualization, and network expansion around known biochemical pathways to predict valuable compound derivations. In line with the standards of open science, we have made the toolbox freely available to the scientific community at http://lcsb-databases.epfl.ch/arbre/. We envision that ARBRE will provide the community with a new computational toolbox and comprehensive search tool to predict and rank pathways towards industrially important aromatic compounds.

## Introduction

The ecological movement towards cleaner and more environmentally friendly science requires the scientific community to revisit current practices in the chemical industry and substitute energy-intensive and toxic syntheses with green chemistry and biotechnological production. Chemical compounds that can be produced with the help of biotechnology include pharmaceuticals, cosmetics, fuels, food additives, and other industrially important substances, many of which are derived from aromatic amino acids. Despite the extreme importance of aromatic compounds, well-characterized pathways for their production are scarce. While some of them can be produced biosynthetically in microorganisms (Huccetogullari et al., 2019), many industrially relevant aromatic compounds are accessed from plant sources or organic synthesis. Thus, there is a growing industrial need for the bioproduction of these high-value molecules.

To produce a desired compound in a microorganism, a biosynthesis pathway, or a sequence of biochemical reactions that generates a target compound from a precursor, must be designed. Computationally predicting such biosynthesis pathways can be done in silico through retrobiosynthesis, in which the target compound serves as a starting point and the biosynthetic pathway is reconstructed backwards to connect the target compound to the desired precursor via a series of biochemical reactions. The importance of retrobiosynthesis for natural and non-natural compounds, as well as the pruning criteria used in retrobiosynthesis, are reviewed in greater detail elsewhere (Hadadi & Hatzimanikatis, 2015)(Jang et al., 2022). Various computational retrobiosynthesis tools have been developed to account for all theoretically possible biosynthesis pathways: BNICE.ch (Hatzimanikatis et al., 2005) (Tokic et al., 2018), RetroPath2.0 (Delépine et al., 2018) (Koch et al., 2019), NovoStoic (Kumar et al., 2018), and ReactPRED (Sivakumar et al., 2016).

BNICE.ch has been used in many biotechnological applications, including the prediction of production pathways for 3-hydroxypropanoate and methyl ethyl ketone precursors (Henry, Broadbelt, et al., 2010)(Tokic et al., 2018). Unlike tools with automatically generated chemical transformation rules (Duigou et al., 2019), BNICE.ch predicts the novel reactions with manually created and generalized enzymatic reaction rules. These expertly curated reaction rules are based on the enzyme commission (EC) classification^1^ and rationally predict novel reactions while preserving the biochemistry. Every reaction predicted with BNICE.ch is annotated with three levels of EC (e.g., 1.1.1.-), which define reaction type and mechanism, but leave the substrate open.

An alternative to directly searching for pathways between the precursors and the target compound is to generate a reaction network that encapsulates biochemistry around the target compound. This approach provides greater flexibility because it allows for exploring the production pathways from a wide set of desired precursors toward an extended set of compounds around the target compound. This concept was presented in the ATLAS of biochemistry series of publications (Hadadi et al., 2016)(Hafner et al., 2020)(Mohammadi- Peyhani et al., 2021). Therein, we used BNICE.ch to predict novel reactions and integrate them with known reactions into a network of all theoretically possible biochemical reactions given a defined basis set of molecules. The reactions are integrated into the network through decomposition of reactions into the reactant-product pairs, which allows to create simple computable graphs (Hafner & Hatzimanikatis, 2021). The first ATLAS of Biochemistry was based on the Kyoto Encyclopedia of Genes and Genomes (KEGG) (Ogata et al., 1999) and was restricted to biological compounds (10K compounds) (Hadadi et al., 2016)(Hafner et al., 2020). Recently, we created ATLASx that covers all known biological and bioactive compounds (further called *biological compounds*, ∼ 1.5 M compounds) and demonstrated that many non-biological organic compounds from chemical compound databases, such as PubChem (Kim et al., 2021) (further called *chemical compounds*) could be generated from biological compounds (Mohammadi-Peyhani et al., 2021). The ATLASx utility in providing biochemical alternatives for complex aromatic compounds was experimentally demonstrated in (Hafner et al., 2021) and (Srinivasan & Smolke, 2021). In this work we endeavored to adapt the ATLASx approach to the specific biochemistry type as it brings several advantages in comparison to using the whole ATLASx network for pathway search.

ATLASx is limited to the compounds in the reference databases as a way to tame the exponential growth of the retrobiosynthesis networks. At the moment, even the comprehensive inclusion of all the compounds in the PubChem database was not yet possible and the PubChem compounds were only included into the ATLASx network as hypothetical products of the biological and bioactive compounds. A potential solution to overcome this issue is to restrict the initial set of molecules based on the specific research or application study, e.g., to build a reaction network around aromatic amino acids metabolism. Limiting the network based on the biochemistry type rather than on the database scope helps to include more relevant compounds into the network while keeping the network size manageable. In this work we generate the comprehensive network around aromatic amino acid biosynthesis which is more than 10 times smaller than ATLASx but includes PubChem compounds in any role, not only as potential products of the biological compounds.

Due to the big size of the ATLASx reaction network, the pathway search algorithm has to enumerate and rank significant number of pathways, which results in the great amount of biosynthetic route alternatives. These biosynthetic route alternatives, while providing a great resource for predicting shortcuts and completely novel routes, might be out of reach for the standard pathway implementation pipelines as they require significant effort in enzyme engineering. In this publication we analyze how restricting the search space to the specific type of chemistry allows to predict pathways that are closer to those existing in nature and therefore are easier to implement.

In addition to finding that the creation of the network centered around specific metabolism type improves the pathway prediction, we saw that additional annotation of the network with the compound properties of interest (e.g., presence of the benzene group in this case) helps to further improve the results of the pathway search.

We report here ARBRE (Aromatic compounds RetroBiosynthesis Repository and Explorer), a computational resource for pathway design centered around key aromatic compounds. ARBRE includes a reactant-product pair network (Hafner & Hatzimanikatis, 2021) that integrates novel reactions predicted using BNICE.ch with known reactions from publicly accessible biochemical databases. To navigate this network, we provide an open-source computational toolbox implemented in python3. The toolbox allows users to search for novel biosynthesis pathways using the NICEpath.ch tool (Hafner & Hatzimanikatis, 2021) by introducing desired source and target compounds. The outputs of pathway searches can be subjected to other computational biochemistry tools, such as BridgIT (Hadadi et al., 2019), and help us identify putative promiscuous enzymes that can catalyze pathway reactions, their associated sequences for direct enzymatic catalysis, or enzymes that can be engineered for novel reaction catalysis. Overall, ARBRE has unprecedented potential for one-click automated, large-scale, systematic analysis of the bioproduction of industrially important aromatic compounds, and it includes a well-curated reaction network accessible for local download and installation. It is a rapid, simple, and user-friendly approach that will complement and improve upon existing methods. We envision that the innovations and features provided by ARBRE will be of broad general interest and applicability for the synthetic biology community. Further the approach in the development of ARBRE could serve as a framework for developing specialized databases around specific chemistries.

## Methods

ARBRE is an open-source resource that allows to automatically combine metrics coming from BNICE.ch, NICEpath.ch and BridgIT to predict and rank the pathway in one click, providing a comparable combined pathway metric for each pathway. ARBRE consists of two main parts: (i) ARBRE repository - the reaction network and data focused around the aromatic amino acid compounds (compounds, reactions, putative enzymes for catalyzing novel reactions, etc.) and (ii) ARBRE toolbox - the computational pipeline to generate pathways from (i) in accordance with user-defined parameters.

### The ARBRE repository

Data for the ARBRE repository (Fig. 1) were generated with BNICE.ch and BridgIT. There are four classes of data: 1) a list of curated compounds used to generate BNICE.ch input; 2) a set of curated known reactions for integration into the RP-pair network; 3) predicted novel reactions based on the enzymatic reaction rules; and 4) novel reactions annotation with the standard Gibbs energy of reaction, conserved atom ratios (CARs) (Hafner & Hatzimanikatis, 2021), and candidate enzymes using BridgIT. From the repository one can extract the following files: network, compound, reaction, and BridgIT prediction files. The network file contains the RP-pairs of the reactions annotated with the CARs. The compound file contains the molecular structures and the canonical names of the molecules in the network, as well as their structural characteristics (see Supplementary Table 1). The reaction file contains information on the stoichiometry and corresponding RP-pairs for each reaction. The BridgIT prediction file contains the top 10 BridgIT predictions for each novel reaction.

**Figure 1.**
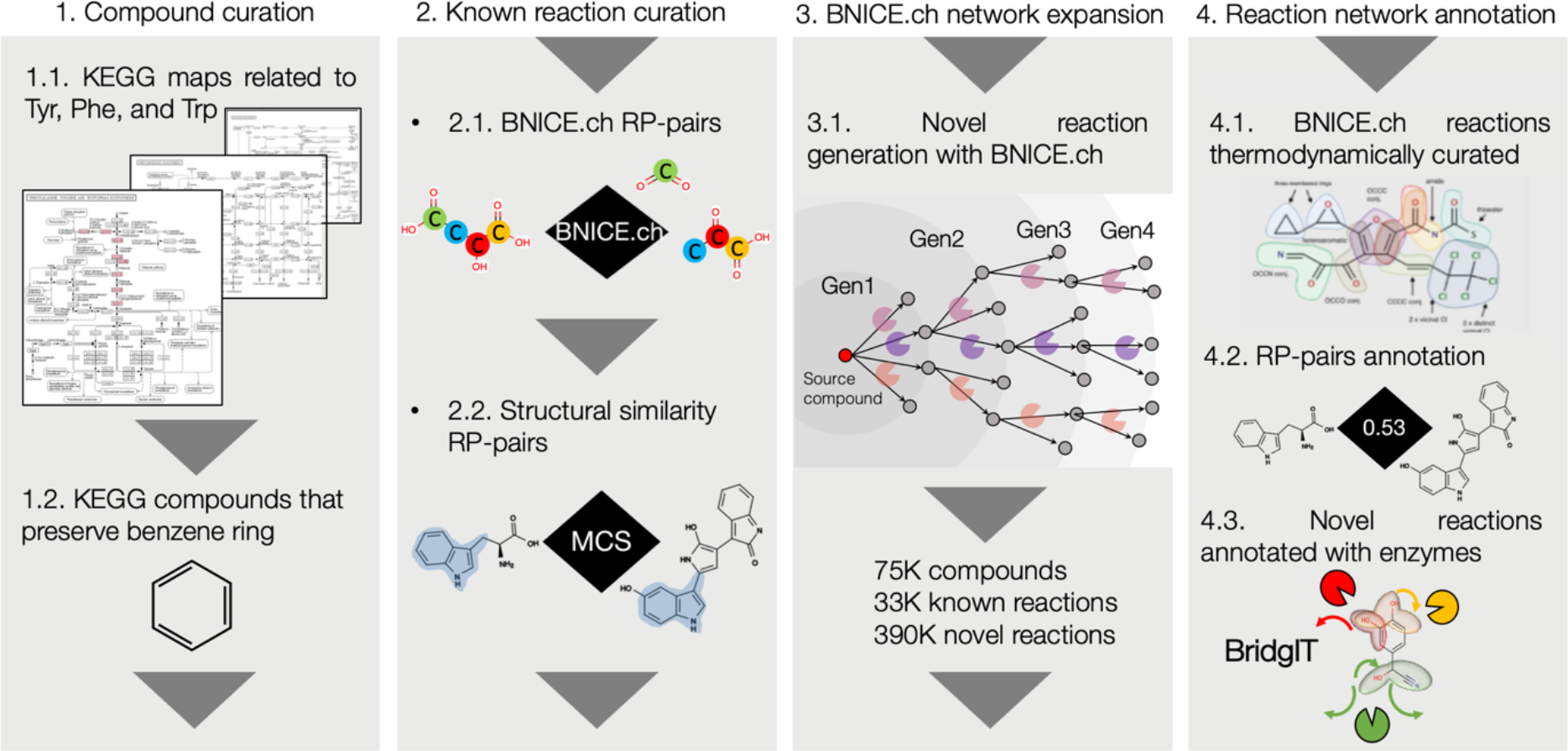
The four steps executed to generate the ARBRE repository around aromatic compounds. MCS – maximum common substructure.

#### Curation of the compound list

The KEGG maps were curated by selecting maps that were solely related to key aromatic amino acid metabolism. KEGG was recursively parsed starting from the “phenylalanine, tyrosine, and tryptophan biosynthesis” map. Compounds and reactions of the encountered maps were collected. Compounds selected as starting points for network generation were sourced from KEGG maps related to “phenylalanine, tyrosine and tryptophan biosynthesis” and had to contain a benzene ring. To maintain consistency within the scope of the study that includes solely small molecule organic compounds synthesis, we excluded polymers (e.g., peptides, glycolipid polymers, and polysaccharides), compounds with over 60 carbon atoms, and compounds without carbon.

#### Curation of the reaction list

The list of known reactions was collected from publicly accessible biochemical databases: MetaCyc (Karp et al., 2002), ModelSEED (Henry, DeJongh, et al., 2010), MetaNetX (Moretti et al., 2021), Brenda(Schomburg et al., 2002), BKMS (Lang et al., 2011), HMR (Pornputtapong et al., 2015), KEGG (Ogata et al., 1999), Reactome (Jassal et al., 2020), Rhea (Alcántara et al., 2012), and BiGG models (Norsigian et al., 2020). The reaction list was curated as described in (Mohammadi-Peyhani et al., 2021). Briefly, the reactions were annotated with BNICE.ch reaction rules if any rule matched a reaction. For reactions that lacked the assigned BNICE.ch reaction rule, the structure-based RP-pairs were assigned based on structural similarity. Finally, the CARs for these pairs were calculated using the maximal common substructure (MCS) algorithm (Raymond & Willett, 2002).

#### Addition of novel reactions

The scope of the reaction network implemented here differs from the previously published version (Mohammadi-Peyhani et al., 2021). To broaden the scope of the reaction network, in addition to reactions with both biological and chemical compounds, we have included reactions between chemical compounds. The BNICE.ch algorithm was applied for four generations (Hadadi & Hatzimanikatis, 2015), starting from 2004 compounds curated from the KEGG maps. In each generation, we added compounds derived in previous generations (including those from PubChem) and reactions including only the PubChem compounds. All compound derivatives with atom conservation lower than 0.34 were excluded. Compounds with over 60 carbon atoms or without carbon were excluded from the network as well.

#### Energy annotation

Known reactions that could be reconstructed with BNICE.ch and novel reactions were assigned a standard Gibbs free energy of reaction based on the group contribution method (Jankowski et al., 2008).

#### RP-pairs annotation

We addressed the challenge to find pathway alternatives with high atom conservation in the retrobiosynthesis reaction network in our recently published tool NICEpath.ch (Hafner & Hatzimanikatis, 2021) by disentangling the complex reaction hypergraph network down to a simple “compound A – compound B” reactant-product pair (RP-pair) network. Tracking the atom conservation of a pair is indispensable for identifying the most conservative biosynthesis pathways. Each RP-pair of the network was assigned a CAR value based on the in-silico atom labeling as described in (Hafner & Hatzimanikatis, 2021). Four distance scores were calculated for every RP-pair in the network based on the CAR:

*- Standard distance* was calculated as 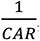, with the value of 1 representing the shortest distance and the value of 1/0.34 (= 2.9) representing the longest accepted distance. The value of 0.34 denotes the standard threshold for the conserved atom ratio proposed in (Hafner & Hatzimanikatis, 2021).

*- Known distance* for associated known reactions was calculated as the standard distance divided by ten, i.e., 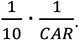. This score was introduced to ensure that known reactions will always be preferred over novel reactions even at the low atom conservation. Specifically, known reactions will have the distance score in the interval (0.1, 0.29) that is always smaller than the distance score of novel reactions ranging from 1 to 2.9.

*- Exponential distance* was calculated as 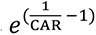

*- Known exponential distance*, similarly to *Known distance*, this score is calculated as 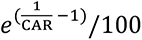 for RP-pairs of known reactions.

#### BridgIT annotation

The novel reactions of the network lack associated enzymes for their catalyzation, which limits their experimental implementation. We applied the BridgIT tool to map these reactions to known biochemistry and find candidate enzymes for their catalyzation. For each reaction, we first kekulized the compounds (Mohammadi-Peyhani et al., 2021) and then generated all possible combinations of kekule forms within the reaction. The resultant kekulized reactions were compared to the BridgIT fingerprint database of the known reactions. The predicted EC numbers that exceeded the similarity score threshold of 0.3 were combined into the unified BridgIT prediction file. The BridgIT prediction file contained the 10 best candidate EC numbers (e.g., EC 1.1.2.1) for each of the reactions based on the BridgIT reaction similarity score. Details about the score calculation and how the threshold was selected are described in (Hadadi et al., 2019).

#### Annotation of compounds with the number of patents and publications

The number of associated patents and publications for each compound provided a quantitative metric on their novelty and whether or not their bioproduction pathways are known. This number was queried from PubChem using the pubchempy python3 library. For each of the compounds, the query was made by defining the INCHIKEY or PubChem ID of the compound. For example, INCHIKEY of norcolaurine is https://pubchem.ncbi.nlm.nih.gov/rest/pug/compound/inchikey/WZRCQWQRFZITDX-AWEZNQCLSA-N/xrefs/PatentID,PubMedID/JSON or its PubChem ID is https://pubchem.ncbi.nlm.nih.gov/rest/pug/compound/cid/440927/xrefs/PatentID,PubMedID/JSON (Kim et al., n.d.). The queries for this study were performed in September 2020.

### The ARBRE toolbox

The ARBRE toolbox (Fig. 2) provides five main functionalities executed in a sequence: 1) finding the pathways within the graph of the ARBRE repository as a sequence of intermediates; 2) assigning reactions to each of the RP-pairs within the initial pathways; 3) assigning potential enzymes to the reactions based on the BridgIT predictions; 4) ranking the pathways according to user-defined metrics; and 5) visualizing the best pathways according to the defined ranking. The inputs of the ARBRE toolbox are network, reaction, compound, and BridgIT prediction files (supplementary table 3), along with user-defined parameters. The output includes the ranked list of pathways and the visualization of the top-ranked pathways.

**Figure 2.**
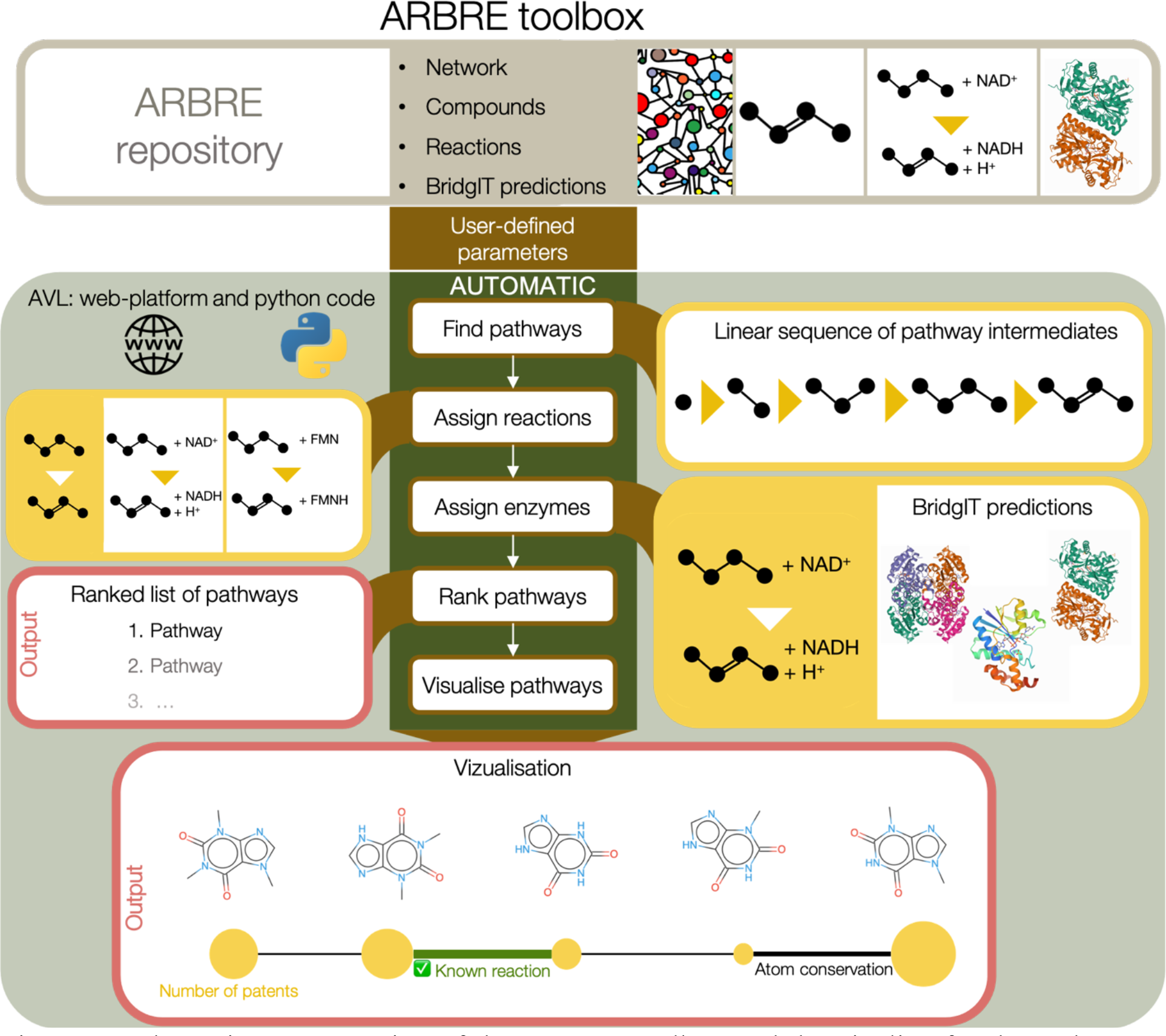
Schematic representation of the ARBRE toolbox and the pipeline for the pathway prediction. The input is the ARBRE resource with the network, compounds, reactions, and BridgIT predictions (see supplementary material for download). The 5 steps of the ARBRE toolbox pipeline (center) are guided by user-defined parameters. Schematic representations of the outputs are shown. Yellow boxes denote intermediate outputs and pink boxes denote final outputs of the pipeline (i.e., ranked list of pathways by score and visualization). In pathway visualization, the circle sizes under the compounds correlate with the associated number of patents. Biotransformations with known enzymes are noted in green, novel in black. The conserved atom ratio (CAR) is reflected in the width of the connection between compounds.

#### Finding pathways based on user-defined parameters

We propose a biosynthesis pathway search that accounts for specific properties of reactions and compounds defined by the end user. This way, the ARBRE network is first filtered in accordance with user-defined compound and reaction parameters, and then the pathway search algorithm generates the initial set of the “raw” pathways within the filtered graph. These pathways, which are represented as a sequence of compound IDs, are then passed to the reaction assignment stage of the workflow.

To allow users to enforce desired properties of the compounds and reactions of the pathway, we introduced adjustable parameters that enable filtering of the network before the pathway search (e.g., after compound filtering, search for the pathways with compounds that fit defined properties).

Compounds can be filtered based on molecular weight, number of atoms, charge, bond and ring structure, atom composition, and biochemical characteristics. Parameters related to the compounds are: (i) number of basic organic atoms (C, O, N, P); (ii) allowed atom types (see full list in Supplementary Table 1); (iii) number of the aromatic and non-aromatic rings, as well as more precise filtering based on ring types (e.g., number of benzene rings). Additionally, undesired compounds can be excluded from the pathway prediction.

Reaction parameters include the atom conservation of the RP-pairs allowed in the pathways; distance transformation, which reflects the importance of the atom conservation for the resulting pathways; whether the known reactions are preferred over novel reactions; and the minimal BridgIT score. The pathway search algorithm accounts for whole-pathway parameters, such as the maximal pathway length and the total number of pathways.

The pathways between any source and target compound can be found with NICEpath search tool (Hafner & Hatzimanikatis, 2021) that is adapted to the ARBRE network. Pathway search is executed using the k-shortest loopless pathways algorithm (Yen, 1971) as implemented with the Python3 NetworkX library (Hagberg et al., 2008). The pathway search algorithm filters for pathways that fall below the defined maximum pathway length and stops if the user-defined number of pathways is reached.

#### Assigning reactions

At this stage, the pathway is represented as a linear sequence of compounds that is translated into reactions. All the associated reactions are assigned for each RP-pair that fit the parameters as defined by the user (e.g., the BridgIT minimal similarity score, preferred known reactions, etc.). The maximum number of the top-fitting reactions assigned to each RP-pair can also be user-defined. If known reactions are absent, then reactions with the best maximum BridgIT score are assigned. Once the reactions are assigned to the RP-pairs, all the possible reaction combinations are generated for each pathway, i.e., we create all possible combinations of cofactors along the pathway. For example, if the pathway “compound A – compound B – compound C” had NAD^+^ - NADH and NADP^+^ - NADPH as possible cofactors for both RP-pairs of the pathway, we would have 4 possible reaction combinations:

1. A+ NAD^+^ = B + NADH → B+ NAD^+^ = C + NADH;
2. A+ NADP^+^ = B + NADPH → B+ NAD^+^ = C + NADH;
3. A+ NAD^+^ = B + NADH → B+ NADP^+^ = C + NADPH;
4. A+ NADP^+^ = B + NADPH → B+ NADP^+^ = C + NADPH.

#### Assigning enzymes

Enzymes that can potentially catalyze reactions of the pathways are assigned based on the BridgIT predictions. By default, the top 3 BridgIT predictions with similarity scores that exceed the user-defined threshold are assigned. However, this parameter can be modified to range from 1 to 10 of the best predictions above the threshold. The standard threshold recommended for considering a BridgIT prediction to be reliable is 0.3 (Hadadi et al., 2019).

#### Ranking pathways

The information about pathways compiled during execution of the first three functionalities of the ARBRE toolbox is used to rank the pathways based on the pathway score. The pathway score accounts for the average BridgIT score (BR, 0 – 1), average atom conservation (CAR, 0 – 1), pathway length (PL, integer value from 1 to the user-defined maximum, where the *shortest PL* is the minimum PL in all pathways), and percent of the known reactions (KR, 0 – 1). By default, the pathway score is calculated according to the formula:

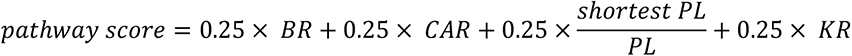

The ideal pathway score is equal to 1. The weighting coefficients for the parameters can be changed in the user-defined parameter file. The results of the pathway ranking are stored withing the pathway file that is the main output of the ARBRE toolbox.

#### Pathway visualization

Pathways with the highest pathway scores are visualized. The ARBRE allows a maximum of 100 top-ranked pathways to be visualized. The 2D structures of the compounds are generated with the rdkit.Chem.Draw package and combined into the pathways with the PIL.Image package. The final image generated for each pathway represents a linear sequence of compounds and computational predications that are ready for user inspection and validation.

#### Preparing the dataset of MetaCyc pathways

The dataset of the MetaCyc pathways was compiled using the dataset of the MetaCyc pathways transformed to linear pathways as described in (Mohammadi-Peyhani et al., 2021). To restrict the analysis to the target space of this article – aromatic compounds derived from chorismate and aromatic amino acids – we excluded the pathways that included compounds without the benzene ring. The resulting dataset included 190 pathways (Supplementary table 4).

#### Comparing the pathway search predictions of ARBRE with ATLASx

To analyze how the results of the step 1 of the ARBRE pipeline (finding pathways based on user-defined parameters) compare with the results of pathway search within the ATLASx network, we performed the pathway search using the same parameters in 4 different network modes: all ARBRE network, ARBRE network with non-aromatic compounds filtered out, and ATLASx network. We used the standard distance and only kept reactant-product pairs with CAR over 0.34. For each pathway we report the rank within each of the 4 networks.

## Results and Discussion

The ARBRE repository was created in four main steps (Fig. 1, Methods): compound curation, reaction curation, BNICE.ch network expansion, and reaction network annotation. The input consisted of 2004 aromatic molecules that were extracted from KEGG maps (Fig. 1, step 1). The reactions of these KEGG maps were annotated with BNICE.ch reaction rules and RP-pairs (Fig. 1, step 2). The resulting network was expanded with novel reactions using BNICE.ch reaction rules (Fig. 1, step 3). All the novel reactions in the final network were annotated with the standard Gibbs energy of reaction, RP-pairs, and BridgIT predictions (Fig. 1, step 4). All the compounds were annotated with structural properties, number of publications, and patents (Methods). The usefulness of the resulting network was illustrated through a set of case studies including the prediction of novel biosynthesis pathways for scoulerine and benzenecarboperoxoic acid and the identification of biosynthetic derivatives of aromatic amino acid and morphine synthesis pathways.

### 2’004 aromatic compounds were extracted from 243 KEGG maps

To define the initial set of target compounds related to aromatic amino acids, we extracted 243 KEGG maps after parsing the KEGG database starting from the “phenylalanine, tyrosine and tryptophan biosynthesis” map. These maps included from 1 to 169 reactions (Supplementary Fig. 1). Out of the 243 KEGG maps, 131 were related to the scope of this study, which includes metabolism, biosynthesis, and biodegradation. The remaining maps were related to classes of biological processes distinct from metabolism, such as signaling pathways. Importantly, all 243 KEGG maps contained aromatic compounds. We extracted compounds and reactions from the 243 KEGG maps, which resulted in a dataset of 6’346 reactions and 2’004 aromatic compounds. Out of these 2’004 compounds, 1’893 could be categorized within the 131 metabolism, biosynthesis, and biodegradation maps. The remaining 111 compounds were solely related to signaling pathways and were not associated with a single metabolic reaction in the remaining 112 KEGG maps.

### Extracted KEGG maps covered 92% of curated reactions

From 6’346 extracted reactions, 3’467 were annotated with a BNICE.ch reaction rule and 2’433 were assigned as chemical structure-based RP-pairs. The remaining 446 reactions fell outside the scope of the study (e.g., polymerization reactions, stereo-isomerization reactions, transport reactions, and reactions involving compounds with undefined structures). Therefore, we ended up with the unified biochemical network of 5’900 reactions, or 92% of the curated reactions.

When we mapped the integrated reactions to their original source in the KEGG maps to see how well our network corresponds to initial information from KEGG, we found that for 114 out of 131 maps over 80% of the reactions were covered in the resulting network. Additionally, 7 maps covered 60 – 80% of the reactions whereas 10 maps covered less than 60% (Fig. 3a, b).

**Figure 3.**
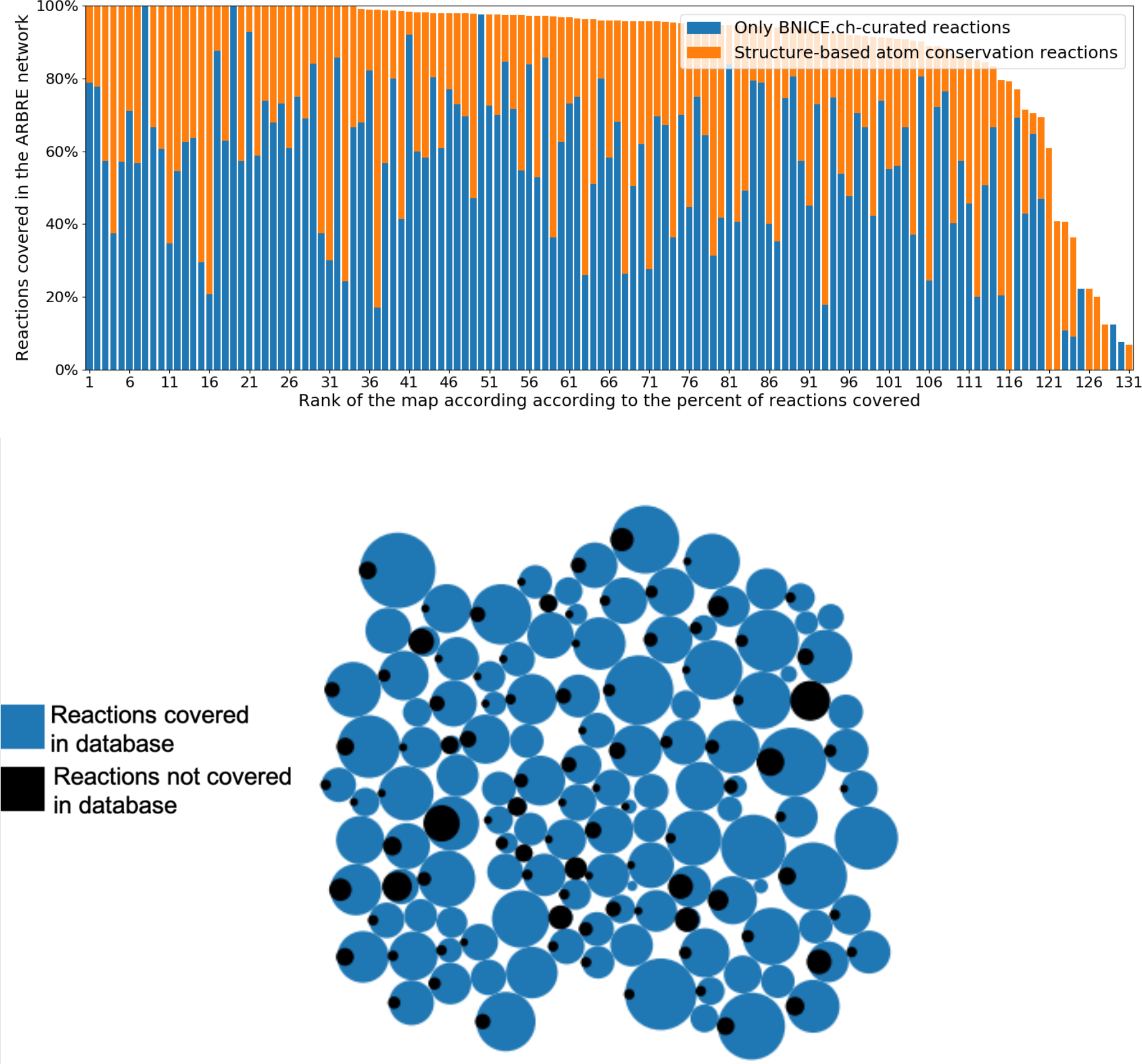
A) KEGG maps ranked by the percent of reactions annotated with BNICE.ch reaction rules (blue) versus reactions with assigned compound RP-pairs included in the network. Reactions beyond the scope of this study are represented by white space. B) Coverage of KEGG maps represented by circles. Circle size corresponds to the total amount of reactions in the map; blue corresponds to number of KEGG reactions covered in the network; and black corresponds to the number of KEGG reactions outside of the network.

The unified RP-pair network included 1’956, or 97%, of all the aromatic compounds from the initial set of KEGG compounds. The remaining 48 compounds belonged to either medicament-related maps without reactions (e.g., bile secretion, vitamin digestion and absorption, and local analgesics) or biosynthesis maps for large molecules outside of the scope of our interests (e.g., porphyrin and chlorophyll metabolism, biosynthesis of type II polyketide products, and biosynthesis of enediyne antibiotics). Taken together, these data demonstrate that the created aromatic amino acids biochemistry network corresponds well to the initial information in the KEGG maps.

### BNICE.ch predicted over 300K novel reactions for generating aromatic compounds

To generate all compounds and reactions that are away by up to 4 reaction steps from the 2,004 extracted aromatic compounds, we applied the BNICE.ch algorithm. The final network integrated 390’175 novel reactions. From the predicted reactions, 65% were transferases, 46% were oxidoreductases, 21% were lyases, 11% were hydrolases, 3.8%were isomerases, and 3.1% were ligases. We found that 0.5% reactions could be performed within the rules of spontaneous enol-ketone transformation or imine formation. From the total set of 63’523 compounds involved in reactions, we found that 303 out of 469 bidirectional generalized enzyme reaction rules of BNICE.ch could localize a reactive site. Additionally, 382’474 of the novel reactions are common to our previous studies on novel reactions generation with ATLASx, while 7’701 reactions are unique to this network. These unique reactions represent hypothetical biotransformations of chemical compounds that are currently not indicated for biological activity, and therefore, only present in the PubChem database. In comparison to previous publications, the ARBRE network includes 1’526 chemical compounds (2% out of 74’053) which were not previously connected to known biochemistry. Overall, these data demonstrate the utility of the ARBRE network for the investigation of potential biosynthesis pathways around aromatic compounds metabolism with greater novelty and predictive power than previously reported datasets.

### The ARBRE network was further enriched with 17K known balanced reactions

After extending the ARBRE network with the novel reactions, we endeavored to include known reactions from various biochemical databases. Enrichment of the network with these known reactions expands the potential integration of pathway search results into computational host organism models. We have provided here two networks: (i) a basic network based on only on BNICE.ch mechanism annotated reactions, and (ii) an extended network with the added known reactions without BNICE.ch mechanism annotation. We found that from the 22’646 known balanced reactions from non-KEGG databases, 5’524 were already predicted during network expansion by BNICE.ch, as described in the previous section. The remaining 17’122 balanced known reactions without BNICE.ch mechanisms were added using structure-based RP-pairs. Overall, we expanded the network to 423’653 reactions with the addition of known balanced reactions from other databases.

### Topological properties of the ARBRE network

We analyzed the network statistics for the basic and the extended networks (Table 1). A total number of 74’053 compounds were included in the extended network with 221’495 RP-pairs. The basic network contained 69’919 compounds and 205’814 RP-pairs.

**Table 1.** ARBRE network statistics. Quantity of compounds, RP-pairs, and connected network components in the network generated around aromatic compounds. The whole network includes all the compounds and RP-pairs without any truncation. Network truncation at CAR = 0.34 includes only connections that are significant for atom conservation along the pathway (see (Hafner & Hatzimanikatis, 2021) for the details). The main network component includes most compounds with the potential for transformation between one another and also includes tyrosine, phenylalanine, and tryptophan.

|  | BNICE-mechanism annotated | All |
| --- | --- | --- |
| Whole network |  |  |
| # compounds | 69,919 | 74,053 |
| # RP-pairs | 205,814 | 221,495 |
| # network components | 241 | 78 |
| Network truncation at CAR = 0.34 |  |  |
| # compounds | 69,189 (99%) | 73,265 (99%) |
| # RP-pairs | 179,908 (87%) | 192,164 (87%) |
| # network components | 1,276 | 790 |
| Only the main network component (part of the network connected to aromatic amino acids) |  |  |
| # compounds | 63,089 (90%) | 70,717 (96%) |
| # RP-pairs | 173,674 (84%) | 190,096 (86%) |
| diameter | 35 | 30 |

We analyzed if the gaps in the network are filled upon the inclusion of known reactions without BNICE.ch annotation. To achieve this we counted the number of components, i.e., subnetworks in which any two compounds belonging to the subnetwork are connected to each other by pathways but these compounds are not connected to any additional compounds in the rest of the network. There were 241 components for 69.9K compounds in the basic network and 78 components for 74.1K compounds in the extended network, meaning that 17K of additional reactions in the extended network significantly improved the network connectivity. The reduced number of components in the extended network benefits the pathway search and expands capabilities towards finding pathways for more source-target compound combinations. Next, we repeated the analysis for the subnetworks with removed RP-pairs with CAR of less than 0.34. These subnetworks had 13% RP-pairs less than the original subnetworks (Table 1). As a result, the two networks had more components, i.e., 1,276 and 790 components for the basic and extended network, respectively. Overall, 99% of the compounds were preserved for each network mode.

To see for how many compounds of the network it is possible to find pathways from aromatic amino acids, we extracted the largest subnetwork containing the key aromatic amino acids – the main component of the network. The compounds from other network components are not connected to the key aromatic amino acids and one cannot find production pathways for them. For the extended network, 96% of all compounds and 86% of all RP-pairs were included into the main component. Thus, almost all compounds of the entire network can be connected to the aromatic amino acid metabolism through biosynthesis. For the main component of the basic network, the longest of all the shortest pathways for all possible pairs of source-target compounds (diameter) was equal to 26. This means, that the longest minimal pathway to transform a source compound into a target compound in the reaction space around aromatic amino acids metabolism has 26 reaction steps.

### Compound structural features analysis for 74K ARBRE compounds

We analyzed compounds for atom composition, ring composition, and other structural properties (i.e., molecular weight, number of rotatable bonds, and charge). We found that 97% of the molecules had from 3 to 40 carbon atoms, 2% had more than 40 carbon atoms, and the remaining 1% had 1 – 2 carbon atoms (Table 2). We also analyzed the number of compounds with the oxygen, nitrogen, phosphorus, sulfur, halogens, and chelated metals (Table 2). In terms of ring structure, 24K compounds had one benzene ring, 11K had two benzene rings, and 1K compounds had three and more benzene rings. In the network, 20K compounds did not have any rings, and the remaining compounds had more complex or non-aromatic ring structures (see Supplementary Table 1 for a list of ring structures). Additionally, 4’916 compounds contained a triple bond. These compound structural properties are provided to users as a filter for pathway search, which will allow to streamline the pathway generation algorithms based on the chemical structure of the compounds.

**Table 2.** Overview of compound atom composition.

| Number of atoms types | Number of compounds | % of total (74K) | Example molecules | Formula |
| --- | --- | --- | --- | --- |
| **Carbon** |  |  |  |  |
| 1–2 | 871 | 1% | carbon dioxide, acetate (C2) | CO<sub>2</sub>, C<sub>2</sub>H<sub>3</sub>O<sub>2</sub> |
| 3–19 | 65,167 | 87% | pyruvate, (C3), (S)-scoulerine (C19) | C<sub>3</sub>H<sub>3</sub>O<sub>3</sub>, C<sub>19</sub>H<sub>21</sub>NO<sub>4</sub> |
| 20–39 | 7,721 | 10% | violacein, (C20), novobiocin (C31) | C<sub>20</sub>H<sub>13</sub>N<sub>3</sub>O<sub>3</sub>, C<sub>31</sub>H<sub>36</sub>N<sub>2</sub>O<sub>11</sub> |
| 40–60 | 1,601 | 2% | paclitaxel, (C47), mithramycin (C52) | C<sub>47</sub>H<sub>51</sub>NO<sub>14</sub>, C<sub>52</sub>H<sub>76</sub>O<sub>24</sub> |
| **Oxygen** |  |  |  |  |
| 1–58 | 70,929 | 94% | benzoate, toluene-4-sulfonate | C<sub>7</sub>H<sub>6</sub>O<sub>2</sub>, C<sub>7</sub>H<sub>8</sub>O<sub>3</sub>S |
| **Nitrogen** |  |  |  |  |
| 1–22 | 41,739 | 55% | L-phenylalanine, L-tryptophan | C<sub>9</sub>H<sub>12</sub>NO<sub>2</sub>, C<sub>11</sub>H<sub>12</sub>N<sub>2</sub>O<sub>2</sub> |
| **Phosphorus** |  |  |  |  |
| 1–8 | 3,968 | 5% | 4-nitrophenyl phosphate, phosphotyrosine | C<sub>6</sub>H<sub>6</sub>NO<sub>6</sub>P, C<sub>9</sub>H<sub>12</sub>NO<sub>6</sub>P |
| **Sulfur** |  |  |  |  |
| 1–9 | 3,723 | 5% | toluene-4-sulfonate, 2-naphthalenesulfonic acid | C<sub>7</sub>H<sub>8</sub>O<sub>3</sub>S, C<sub>10</sub>H<sub>8</sub>O<sub>3</sub>S |
| **Halogens (Cl, Br, I, F)** |  |  |  |  |
| 1–10 | 4,269 | 6% | chlorobenzene, chlorothalonil | C<sub>6</sub>H<sub>5</sub>Cl, C<sub>8</sub>Cl<sub>4</sub>N<sub>2</sub> |
| **Chelated metals (Mg, Fe, Co, Cu, Ni or Mo)** |  |  |  |  |
| 1–7 | 138 | 0.2% | heme, chlorophyll a | C<sub>34</sub>H<sub>30</sub>FeN<sub>4</sub>O<sub>4</sub>, C<sub>55</sub>H<sub>75</sub>MgN<sub>4</sub>O<sub>5</sub> |

### The ARBRE network contains 34K unpatented compounds associated to key aromatic **acids**

The number of patents that are assigned to a compound is a factor in the consideration of potential therapeutic substances in the pharmaceutical industry. We first evaluated the number of patents assigned to the initial set of compounds obtained from the KEGG maps and found that 301 out of 2’004 compounds (15%) lacked patents (Fig. 4a). We then evaluated the number of compounds in the predicted network with patents. The extended network comprised 74’053 compounds where 34’080, or 46%, were not yet patented (Fig. 4b). These compounds may have potential as novel targets for biotechnological production.

**Figure 4.**
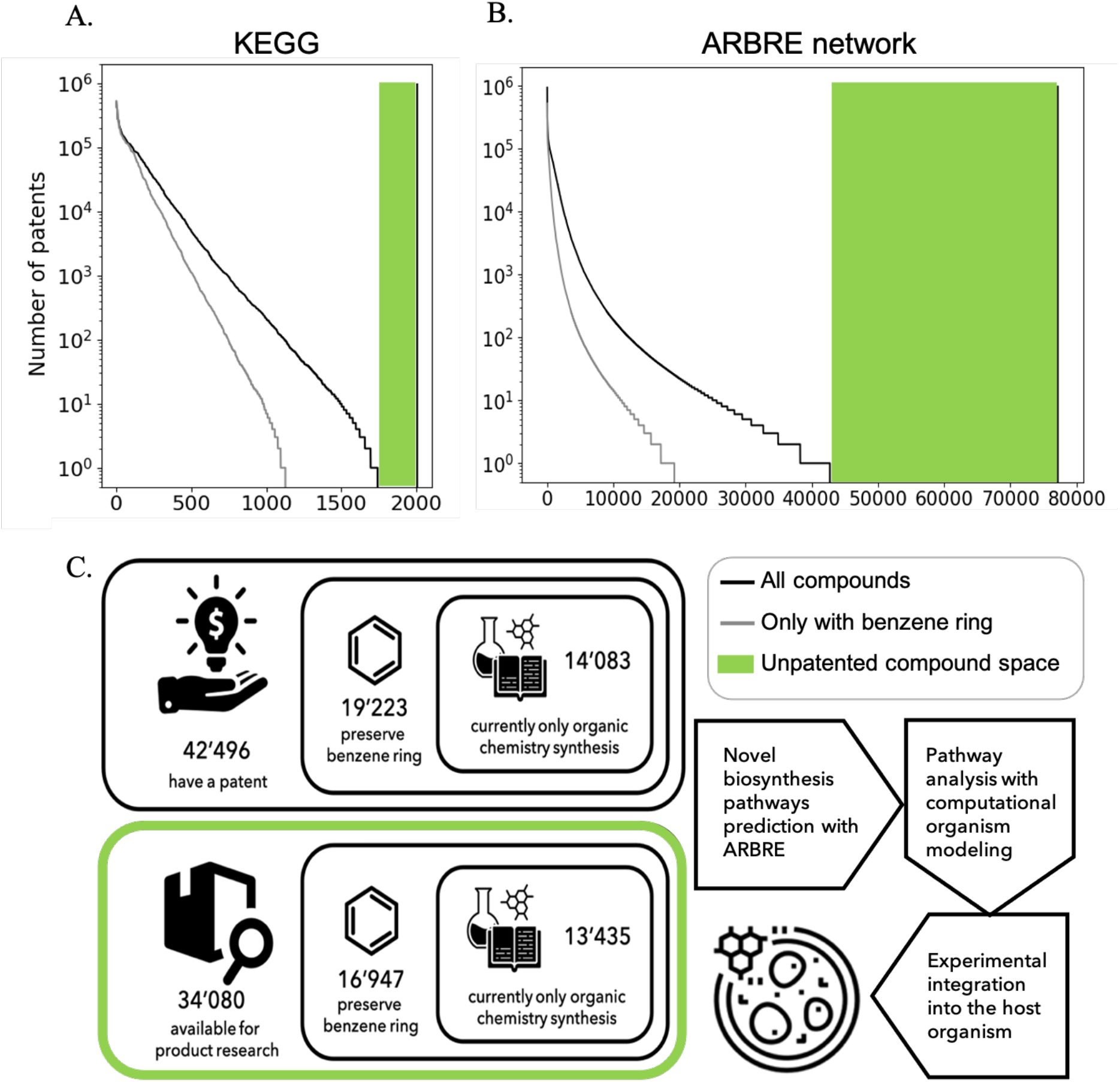
Number of patents per compound. Black line on the right indicates total number of compounds. A) Number of patents per compound in the initial dataset of 2,004 compounds. B) Number of patents per compound in the extended network. C) Schematic representation of how the possibilities for the discovery and patenting of aromatic compounds and their biosynthesis pathways are to be assessed.

### ARBRE assigns a biochemical reaction to 55K PubChem compounds

The search for biosynthetic alternatives for synthetic compounds that lack biosynthesis routes is attractive for biotechnological innovation. Novel biodegradation routes for synthetic chemicals can provide alternatives to more toxic chemical processes. Out of the 74K compounds integrated into the extended network, 55’589 of these compounds originating from PubChem were not present in any biological or bioactive database. This means that it is theoretically possible to integrate these compounds into the metabolism. From this subset of compounds, 27’518 have at least one benzene ring in the structure, which suggests the possibility of biochemically catalyzed pathway predictions starting from aromatic amino acids (Fig. 4c). Selecting these compounds as potential biosynthesis targets can lead to a greener chemistry and new patents for biosynthesis of valuable chemicals.

### Native pathways for aromatic compounds are more likely to be found in the ARBRE network than in the ATLASx network

To test ARBRE network as a basis for pathway predictions, we compared the rank of the native pathway according to the step 1 of ARBRE with the rank of the native pathway within the ATLASx. ATLASx network includes over 5M reactant-product pairs and is more than 10 times bigger than ARBRE network. For the comparison we used the dataset of 190 pathways of aromatic compounds coming from MetaCyc (see Methods). Both ATLASx and ARBRE networks covered all the pathways in the dataset, however, differences in the rank of the native pathways were observed. Our hypothesis was that since ARBRE is centered around aromatic amino acids it predicts the native aromatic pathways with better rank than ATLASx network as it avoids shortcuts in pathways through alternative chemicals.

We observed that the native MetaCyc pathways were ranked higher in the ARBRE network than in the ATLASx network confirming our hypothesis (Fig. 5). The improvements in rank were observed for 95 pathways out of 190. On average, pathways were ranked 124 points better within ARBRE than in the ATLASx. In the extreme case, for the pathway PWY-7256 the ATLASx ranked the native pathway 5833 positions lower than ARBRE network. The filtration of the ARBRE network to keep only the molecules with benzene ring helped to further improve the ranking of the native pathway 9 points on average in comparison to the unfiltered ARBRE network and up to 810 points better for the pathway PWY-1121 (rank 994 for the unfiltered vs 184 for the filtered ARBRE network). These results demonstrate that creation of the specialized networks and their curation can significantly improve the quality of pathway prediction for the specific metabolism type.

**Figure 5.**
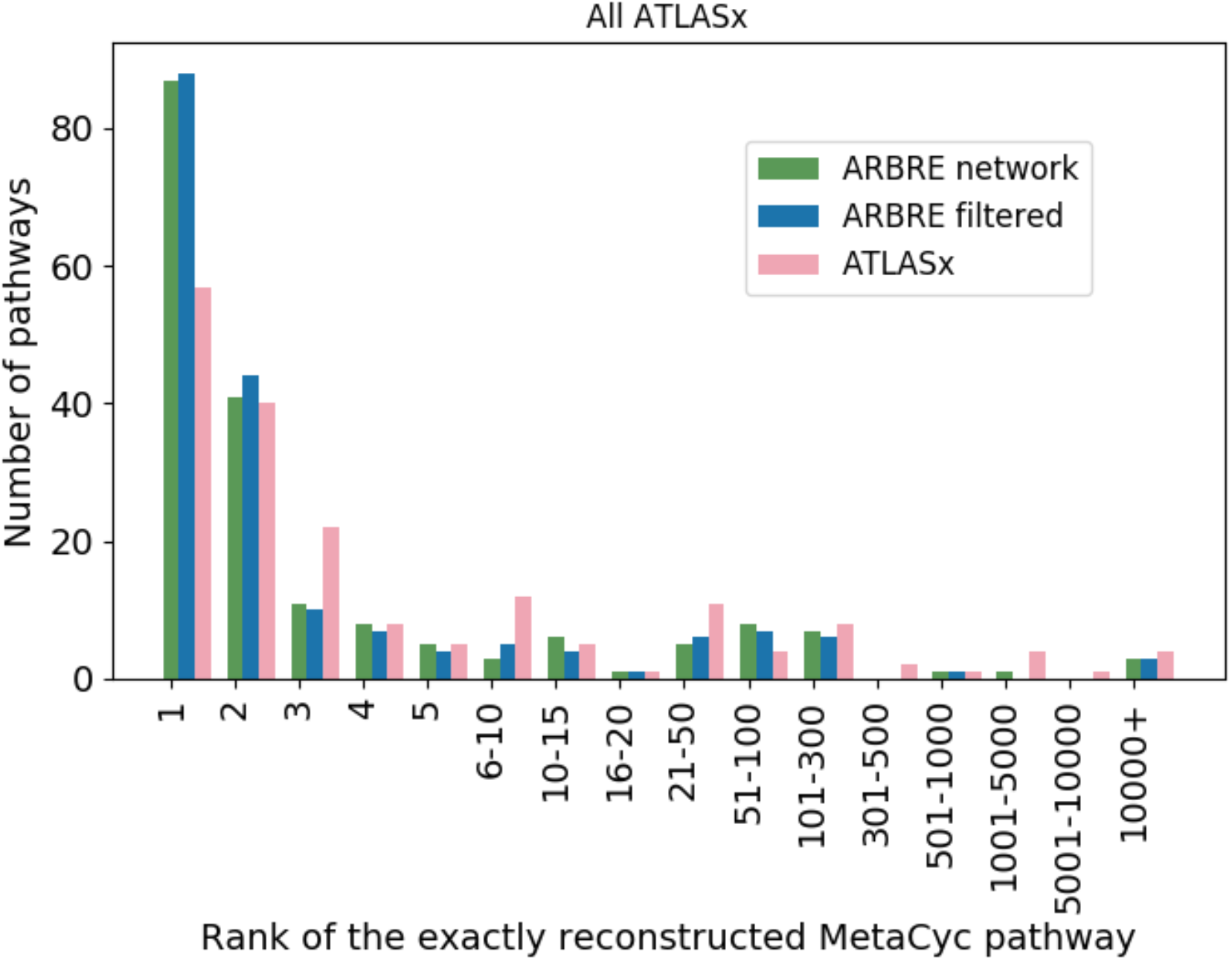
Rank of the native pathway according to the pathway search algorithm performed within the network of ARBRE network, ARBRE network filtered to only aromatic compounds, and ATLASx network.

### ARBRE can resolve common issues in pathway reconstruction

In this section, we discuss the user-defined parameters and how they can affect the results of the pathway search algorithm.

- Minimizing enzyme engineering work

A major concern in synthetic biology for creating new bioproduction or biodegradation pathways for compounds is the effort required to create enzymes capable of catalyzing the computationally predicted reactions. ARBRE allows users to give preference to reactions catalyzed by known enzymes. Alternatively, users can use the default pathway search algorithm settings to optimize the number of pathway steps and the final yield. As an example, we used the pathway search algorithm to find production pathways from tyrosine towards the natural compound umbelliferone, which is used in commercial cosmetic products (Fig. 6). The highest-ranked pathway consisted of four steps, with the last two steps requiring enzyme engineering (Fig. 6, left). The alternative search with a preference for known reactions returned a five-step pathway, with only one step requiring enzyme engineering (Fig. 6, right). While this pathway has more reaction steps, its implementation is likely less time-consuming because only one step needs enzyme engineering.

- Optimizing the total atom conservation of the pathway

**Figure 6.**
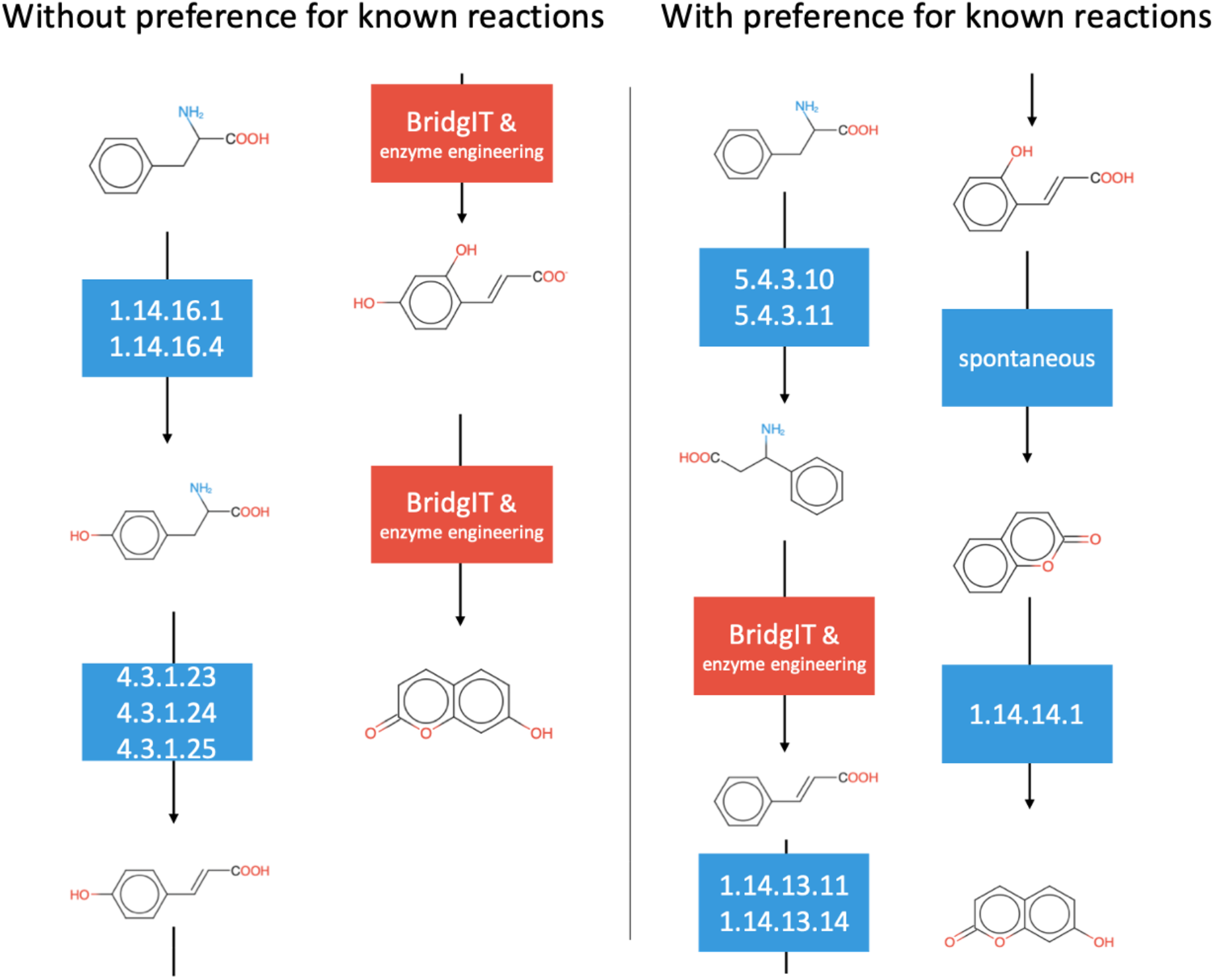
Difference in pathway search results to umbelliferone from tyrosine depending on the preference for known reactions. Blue boxes: steps with a known enzyme with indicated EC class). Red boxes: steps with unknown enzyme where BridgIT results are applied to find a candidate enzyme for their catalyzation.

In nature, longer pathways are sometimes advantageous as they are more energetically feasible. For example, natural biosynthesis pathways of secondary metabolites derived from the key aromatic amino acids can often exceed 10 steps, e.g., biosynthesis of isoquinoline alkaloids papaverine (Han et al., 2010) and colchicine (Nett et al., 2020) from tyrosine. However, computational predictions sometimes produce shortcuts in such naturally long pathways because the algorithms try to minimize the number of the steps even though atom conservation is compromised in the resulting routes. In order to address this issue, in ARBRE we use the exponential transformation of distance as a parameter in the pathway search (Methods). Using exponential distance can also allow to select CAR threshold lower than recommended 0.34. To illustrate this feature, we analyzed pathways from tyrosine to papaverine (Fig. 7). In both searches the CAR threshold as set to 0.25. We observed that the shortest predicted pathway with the standard distance lost the atoms of the initial molecules by making a synthetic “shortcut” via hydrogen peroxide cofactor (Fig. 7, right). In contrast, the pathway predicted with exponential distance had more reaction steps but conserved the atoms of the source molecule (Fig. 7, left). This way, we obtain the more reliable predictions with the exponential distance because this type of the distance transformation exacerbates the difference between low and high atom conservation connections. As such, the exponential distance should be used in the pathway search if there is a need of maximizing the atom conservation to improve the yield of the predicted pathways.

**Figure 7.**
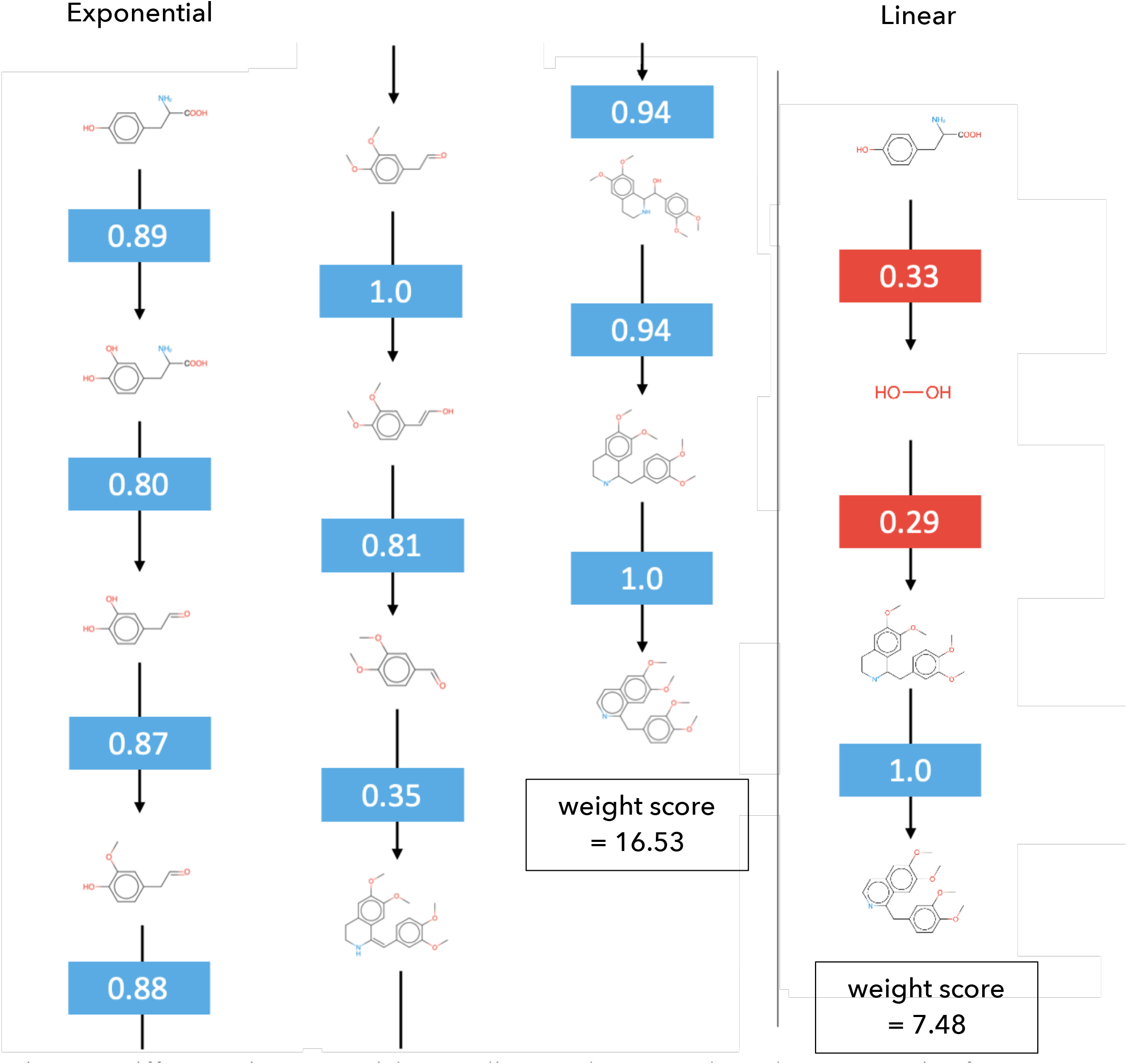
Differences in exponential versus linear pathway search results to papaverine from tyrosine. CARs for each step are indicated. Blue: CARs that are allowed by default. Red: CARs below the default threshold.

### Case studies

To illustrate potential applications of ARBRE, we performed a set of case studies where we used this resource to (1) predict biosynthesis/biodegradation pathways for a biological and a chemical compound, and (2) expand the space of potentially derivable compounds around the existing pathways.

- Case study 1a: biosynthesis pathways toward scoulerine

We used ARBRE to predict biosynthesis pathways toward scoulerine, an important intermediate in the biosynthesis of isoquinoline alkaloids. The predicted pathways were ranked according to the pathway score (Methods). The top-ranked pathway contained 7 reaction steps (Fig. 8a). The average atom conservation was 0.89, the pathway did not have novel reaction steps and the total pathway score was 0.96. This pathway started with tyrosine as a precursor and followed through 3,4-dihydroxy-phenylalanine to dopamine. Using norcoclaurine synthase, dopamine was further condensed with 4-hydroxyohenylaldehyde to form norlaudanosoline, which was downstream decorated with small chemical groups to form reticuline, 6-O-methylnorlaudanosoline, 3-hydroxy-N-methylcoclaurine, and finally scoulerine. The predicted pathway corresponds to known routes to isoquinoline alkaloids that require norcoclaurine synthase to form the structural backbone (as reported in KEGG, https://www.kegg.jp/pathway/map00950).

- Case study 1b: biosynthesis pathways toward benzenecarboperoxoic acid

**Figure 8.**
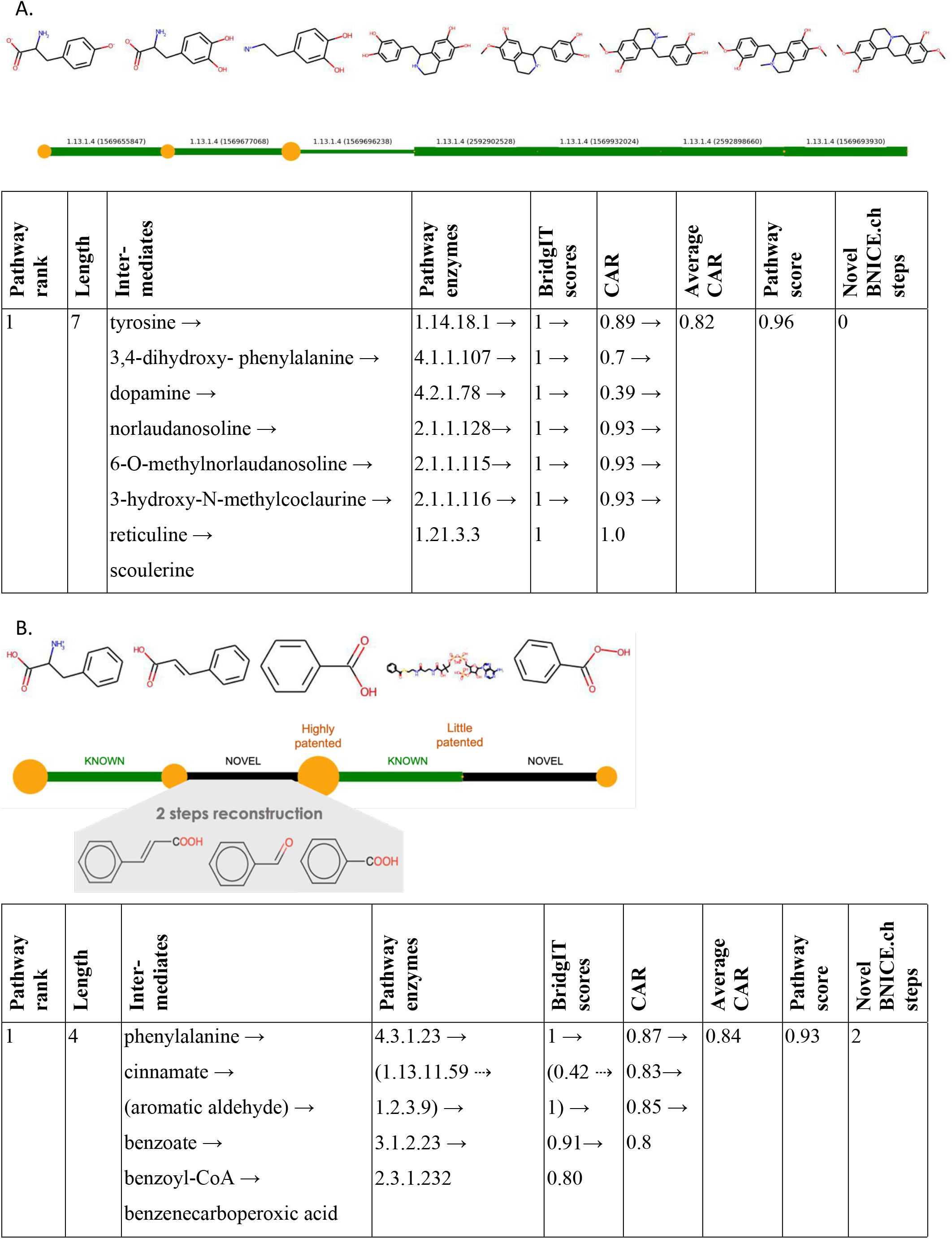
Example pathway annotation for prediction. Size of the yellow circle corresponds to the number of associated patents. Green lines correspond to reactions in public databases. Black lines correspond to novel BNICE.ch reactions. CAR = conserved atom ratio. A) Predicted biosynthesis pathway for scoulerine (PubChem ID: 439654, LCSB ID: 1467886249) from tyrosine (PubChem ID: 6057, LCSB ID: 1467866617). B) Predicted biosynthesis pathway for benzenecarboperoxoic acid (PubChem ID: 523077, LCSB ID: 6154966) from phenylalanine (PubChem ID: 6140, LCSB ID: 1467866580). One the right: two-step reconstruction of the known multistep reaction without enzyme assigned (through web- interface).

We selected benzenecarboperoxoic acid as a proof-of-concept to demonstrate novel biosynthesis pathway prediction for patented compounds that have characterized only organic chemistry synthesis routes. Benzenecarboperoxoic acid is a compound with over 100K patents. With ARBRE, we found pathways to transform phenylalanine into benzenecarboperoxoic acid through enzymatic transformations (Fig. 8b). The top ranked pathway included just four reaction steps, where two reaction steps were associated with known enzymes while other two reaction steps were novel. The novel transformation of cinnamate into benzoate that could not be assigned an enzyme based on BridgIT predictions was reconstructed in 2 reaction steps (Fig. 8b). The average atom conservation was 0.8, which suggests high yield from the resulting pathway. Before ARBRE, such biosynthesis pathways toward chemical compounds could not be found from common databases of predefined pathways based on known biochemical reactions.

The same algorithm can be applied to any of the compounds within the ARBRE network to predict potential biosynthesis pathways. The search for biodegradation pathways can be performed by switching target and source compounds.

- Case study 2a: expanding the space of potentially derivable compounds around aromatic amino acid synthesis

Existing native and heterologous biosynthesis pathways can benefit from expansion to new industrially valuable derivatives. We propose an approach to analyze the ARBRE network in search of potential high-value derivatives within biochemical proximity of the characterized biosynthesis pathways. We used the number of patents or publications related to high-value derivatives as a search criterion. Such approach was successfully used for the expansion of heterologous biosynthetic pathways toward noscapine (Hafner et al., 2021) and tropane alkaloids (Srinivasan & Smolke, 2021) to their potential derivatives. This functionality is now implemented in ARBRE.

Here, we demonstrate the reconstruction and enrichment of the native pathways of the three main aromatic acids tyrosine, phenylalanine, and tryptophan from shikimate. There is a variety of compound derivatives for aromatic amino acids biosynthesis (Fig. 9a). Most patented aromatic derivatives of the pathway were found to be cinnamate, 4-hydroxybenzoate, phenylacetic acid, hydroquinone, and aniline. The most highly patented compounds are centered around tyrosine and phenylalanine. Conversely, most compounds derived from tryptophan have few associated patents. A total of 1,800 compounds were within biochemical proximity of aromatic amino acids biosynthesis (with the atom conservation threshold set to 0.34).

- Case study 2b: expanding the space of potentially derivable compounds around morphine synthesis

**Figure 9.**
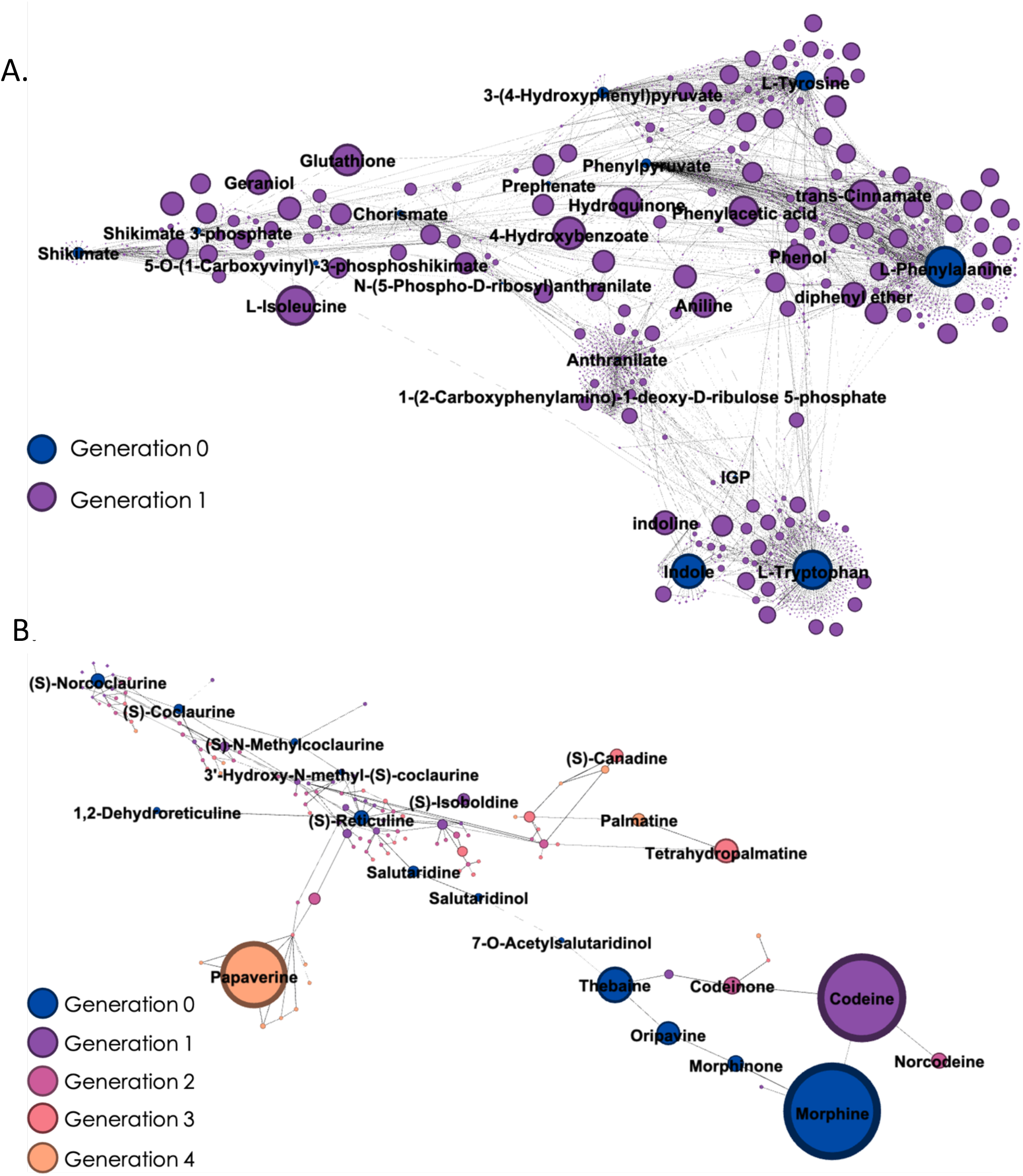
Known pathways enrichment. Number of patents associated with each compound as represented by circle size. A) Enrichment of tyrosine, phenylalanine, and tryptophan biosynthesis pathways from shikimate with potential biosynthetic derivatives 1 reaction step away (minimum CAR = 0.34). The top 15 compounds are determined by the number of associated patents are labeled. B) Enrichment of the morphine biosynthesis pathway from norcoclaurine with biosynthetic derivatives 4 reaction steps away (minimum CAR = 0.8). The top 8 patented compounds are named (codeine, papaverine, etc).

To demonstrate the ability of the ARBRE network to predict complex pathways and their derivatives, we selected the morphine biosynthesis pathway from the norcoclaurine (Fig. 8b), one of the longest isoquinoline alkaloid biosynthesis pathways. First, we verified that the ARBRE network included the original morphine biosynthesis pathway. We expanded this biosynthetic network with all potential derivatives that were 4 reaction steps away. We set a high atom conservation to select for the highest yield of these complex compounds. In the process, we found another analgesic, codeine, with equal amounts of associated patents as morphine that is one reaction step away. We also found other therapeutic compounds that were separated from morphine: isoboldine (antihypertensive and psychoactive) derived from reticuline, one reaction step away; codeinone and norcodeine (analgesics), 2 reaction steps away; canadine (antioxidant) and tetrohydropalmatine (analgesic), 3 reaction steps away; and palmatine (antihypertensive and anti-inflammatory) and papaverine (antispasmoic), 4 reaction steps away. These results demonstrate that ARBRE network can be successfully applied to select compounds for biosynthesis based on previously characterized pathways and expand the scope of applications of the existing pathways.

## Conclusion

We have generated a computational resource, called ARBRE: Aromatic compounds RetroBiosynthesis Repository and Explorer, which is the first open-source code implementation of the BNICE.ch-based retrobiosynthesis pipeline. ARBRE is a novel and comprehensive network of known and predicted biochemical reactions centered around aromatic amino acid metabolism and its derivatives that can be data-mined and analyzed using a suite of computational tools for pathway prediction. Users can evaluate the resulting pathways and assign enzymes to the reactions of interest from the provided BridgIT predictions. Extensive patent analysis of the ARBRE network compounds suggests avenues for new alternatives for product engineering and opportunities to replace chemical synthesis with biotechnological synthesis. We envision that ARBRE will become an indispensable tool for researchers interested in finding alternative biosynthesis or biodegradation routes for aromatic pharmaceuticals, small-molecule nutrients, and other bioagents for which no known biochemical routes exist. The proposed implementation can be further extended to larger reaction networks with an increased number of compounds that can be synthesized biochemically.

## Acknowledgements

We thank Dr. K. Butler and Dr. L. Miskovic for valuable feedback in the preparation of the manuscript. Funding for this work was provided by the Swiss National Science Foundation (SNSF grant agreement 200021_188623 and NCCR Microbiomes grant agreement 51NF40_180575), the European Union’s Horizon 2020 research and innovation program under grant agreement 814408 and Marie Skłodowska-Curie grant agreement No 72228, and the Ecole Polytechnique Fédérale de Lausanne (EPFL).

---

**Supplementary Figure 1.**
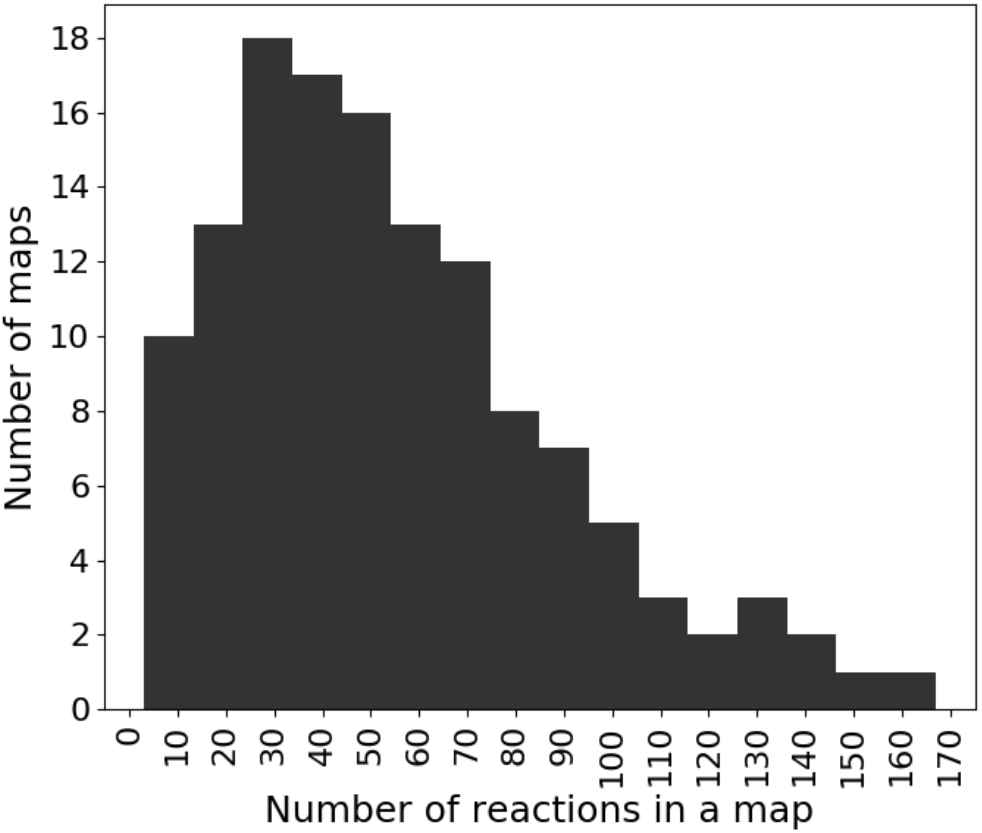
Distribution of number of reactions per KEGG map in the initial dataset (131 KEGG maps related to metabolism, biosynthesis, or biodegradation).

**Supplementary Figure 2.**
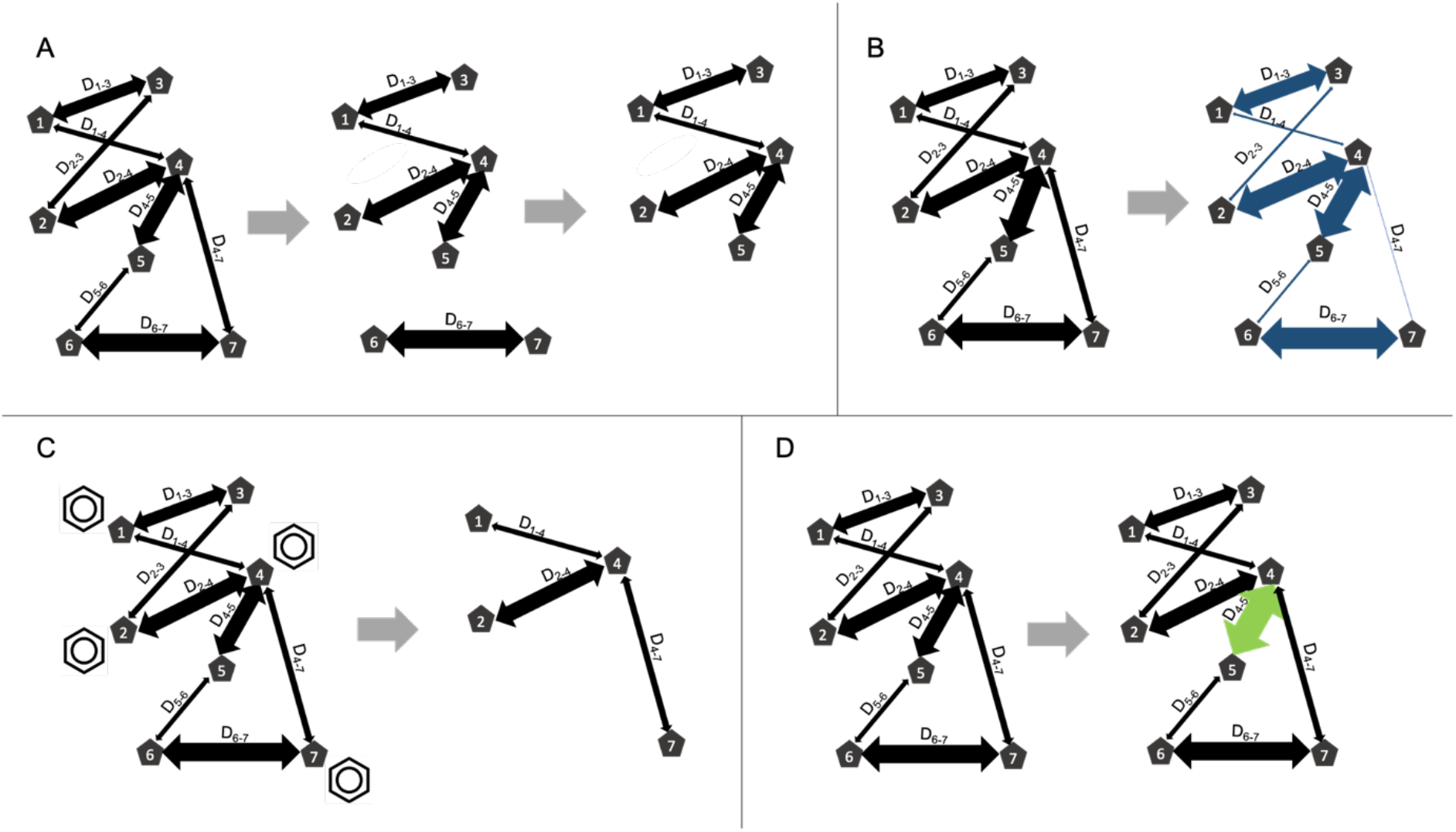
Types of network transformations that are applied to the reaction network for pathway search according to user-defined parameters. A) Removal of RP-pairs with the atom conservation below the defined threshold. B) Transforming of distances to force the algorithm to select pathways with higher atom conservation at the cost of pathway length (exponential transformation). C) Exclusion of compounds that lack the requested substructure. D) Transformation of the distances for pathways with known reactions steps.

**Supplementary Table 1.** Structural properties of compounds available for filtering

|  |  |
| --- | --- |
| Size | Number of atoms, MW |
| Bounds structure and rings | Number of rings, number of rotatable bonds, triple bond presence |
| Rings type count | benzene, pyridine, pyrimidine, pyridazine, imidazole, imidazole cation, pyrrole, pyrazine, furan, pyrazole, thiophene, isoxazole, isoxazole cation, oxazole, thiazole, thiazole cation, 1,2,3,4-tetrazole, 1,2,3,5-tetrazole, 1,3,4-triazole, 1,2,3-triazole, 1,2,4-triazole, 1,3,4-thiadiazole, 1,2,3-thiadiazole cation, other ring type |
| Atom types count | H, Cu, Co, Mg, I, F, C, Cl, B, Se, As, Na, Ni, Mn, Ca, Br, N, Cr, Cd, Mo, K, Ag, Hg, Fe, O, P, S, Si |

**Supplementary Table 2.**
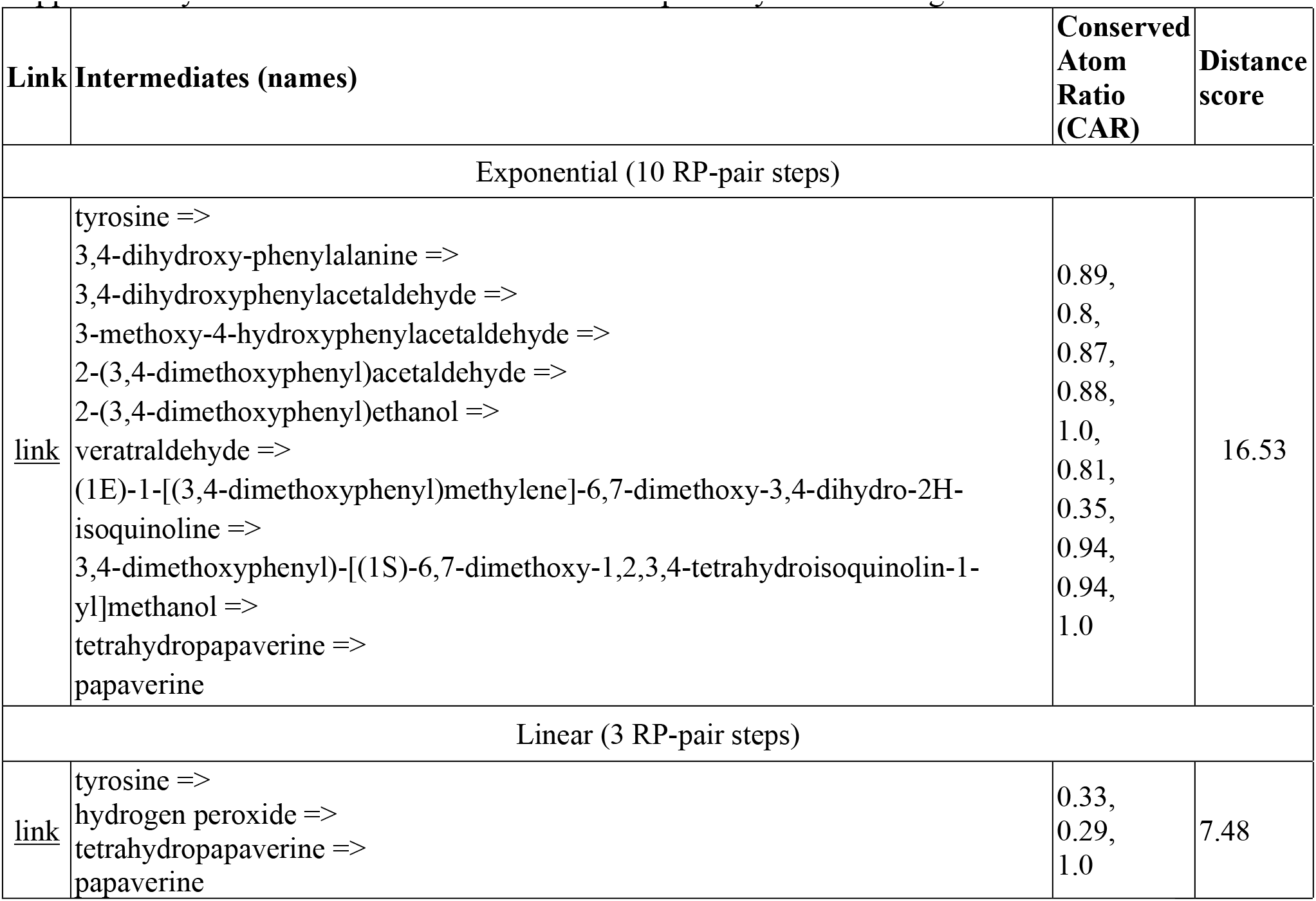
Detailed annotation of the pathways from the fig. 7 Top ranked pathways to papaverine from tyrosine according to the pathway search algorithm in the exponential and linear distance mode and with the rest of the parameters equal.

| Link | Intermediates (names) | Conserved Atom Ratio (CAR) | Distance score |
| --- | --- | --- | --- |
| Exponential (10 RP-pair steps) |  |  |  |
| link | tyrosine =><br>3,4-dihydroxy-phenylalanine =><br>3,4-dihydroxyphenylacetaldehyde =><br>3-methoxy-4-hydroxyphenylacetaldehyde =><br>2-(3,4-dimethoxyphenyl)acetaldehyde =><br>2-(3,4-dimethoxyphenyl)ethanol =><br>veratraldehyde =><br>(1E)-1-[(3,4-dimethoxyphenyl)methylene]-6,7-dimethoxy-3,4-dihydro-2H-isoquinoline =><br>3,4-dimethoxyphenyl-[(1S)-6,7-dimethoxy-1,2,3,4-tetrahydroisoquinolin-1-yl]methanol =><br>tetrahydropapaverine =><br>papaverine | 0.89,<br>0.8,<br>0.87,<br>0.88,<br>1.0,<br>0.81,<br>0.35,<br>0.94,<br>0.94,<br>1.0 | 16.53 |
| Linear (3 RP-pair steps) |  |  |  |
| link | tyrosine =><br>hydrogen peroxide =><br>tetrahydropapaverine =><br>papaverine | 0.33,<br>0.29,<br>1.0 | 7.48 |

## Footnotes

1 “Enzyme nomenclature: Recommendations (1992) of the Nomenclature Committee of the International Union of Biochemistry and Molecular Biology. Pp 862. Academic Press, San Diego. 1992 ISBN 0-12-227165-3,” 1993

